# TNFR2 induced priming of NLRP3-inflammasome via RIPK1 leads to pyroptosis in XIAP deficient cells

**DOI:** 10.1101/550749

**Authors:** Janin Knop, Lisanne M. Spilgies, Stefanie Rufli, Ramona Reinhart, Lazaros Vasilikos, Monica Yabal, Philipp J. Jost, Harald Wajant, Mark D. Robinson, Thomas Kaufmann, W. Wei-Lynn Wong

## Abstract

Recent data suggests that LPS stimulation can trigger inflammasome activation through a TNFR2/TNF/TNFR1 mediated loop in *xiap*^−/−^ macrophages. Yet, the direct role TNFR2-specific activation plays in the absence of XIAP is unknown. We found TNFR2-specific activation lead to cell death in *xiap*^−/−^ myeloid cells, particularly in the absence the RING domain. RIPK1/TAK1 kinase activity downstream of TNFR2 resulted in a TNF/TNFR1 cell death independent of necroptosis. TNFR2-specific activation lead to a similar inflammatory NF-kB driven transcriptional profile as TNFR1 activation with the exception of up-regulation of NLRP3 and caspase-11. Activation and up-regulation of the canonical inflammasome was mediated by RIPK1 kinase activity and ROS production. While both RIPK1 kinase activity and ROS production reduced cell death as well as release of IL-1β, the release of IL-18 was not reduced to basal levels. This study supports, targeting TNFR2 specifically to reduce IL-18 release in XIAP deficient (XLP-2) patients.

## Introduction

Full length tumor necrosis factor (TNF) is a membrane bound protein, where the extracellular domain can be cleaved by TNF converting enzyme (TACE) to release a soluble form(Bell et al., 2007). Soluble and membrane TNF can bind and activate TNF receptor 1 (TNFR1) while only the membrane bound form triggers signaling from TNFR2 (Wicovsky et al., 2009).

The effect of TNFR1 activation has been intensely studied. The outcome of TNF/TNFR1 signaling can range from production of other cytokines, proliferation, survival and differentiation as a result of downstream activation of NF-κB and MAP kinases. While activation of TNFR1 does not normally lead to cell death, the capacity for TNFR1 to induce apoptosis or necroptosis is swayed by the ubiquitylation and phosphorylation of RIPK1 and the activation of pro-survival signals mediated by NF-κB and MAPK pathways(Dondelinger et al., 2017; Jaco et al., 2017; Menon et al., 2017). These survival signaling pathways inhibit the activation of complex II, and the possibility of apoptosis through caspase-8 activity. In the absence or inhibition of caspase-8 activity, necroptosis ensues through RIPK1 kinase activity, RIPK3 and MLKL necrosome activity(Newton et al., 2014; 2016). Phosphorylation of MLKL causes a conformational change, allowing for pore formation and the release of intracellular components as well as damage-associated molecular patterns (DAMPs)(Murphy et al., 2013).

The expression of TNFR2 on cells of the immune system and endothelial cells is highly regulated. Upon binding of membrane-bound TNF to TNFR2 (Grell et al., 1995) a complex is formed that consists of TRAF2, cIAP1, cIAP2, TRAF3 (Rothe et al., 1995). The activation leads to the degradation of cIAP1 and TRAF2 and signals through the non-canonical NF-κB pathway(Rauert et al., 2010). Due to the absence of a death domain, TNFR2 is considered to be mainly involved in survival and maturation of immune cells. Previous data suggest that in some tumor cell lines, TNFR2 can regulate cell death by the loss of cIAP1 and TRAF2 and production of TNF thereby TNFR1 activation(Grell et al., 1999). In addition, immortalized macrophages died in response to TNFR2 via necroptosis when caspases were inhibited(Siegmund et al., 2018; 2016).

In response to LPS, XIAP deficient myeloid cells undergo necroptosis dependent on RIPK3, mediated by TNF(Lawlor et al., 2015; Wicki et al., 2016; Yabal et al., 2014). The loss of XIAP pre-disposes myeloid cells to LPS-induced sensitivity allowing for inflammasome activation(Lawlor et al., 2015; Wicki et al., 2016; Yabal et al., 2014). The activation of the inflammasome, which can be led by NLRP3, ASC and pro-caspase-1 or non-canonical pathway caspase-11, leading to pyroptosis. The stimulation of TLRs primes the cell for inflammasome activation through the upregulation of inflammasome components such as NLRP3(Bauernfeind et al., 2009; Martinon et al., 2002). A secondary stimulus is required to activate the inflammasome *in vitro*. When caspase-1 is cleaved and thereby activated, it will cleave its downstream targets IL-1β and gasdermin D. Cleaved gasdermin D will eventually cause pore formation that facilitates the release of cleaved IL-1β, IL-18 and the osmotic lysis of the cell(Kayagaki et al., 2015; Meunier et al., 2014). XIAP deficient myeloid cells show increased inflammasome activation without any additional stimuli traditionally needed *in vitro*. The inflammasome activation is dependent on RIPK3/caspase-8 activation driven by TNF/TNFR1/TNFR2 stimulation in response to LPS(Lawlor et al., 2015; 2017; Yabal et al., 2014). These data suggest that the use of anti-TNF compounds may benefit patients with loss of function mutations in X-linked inhibitor of apoptosis (XIAP). Often referred to as X-linked lymphoproliferative syndrome (XLP-2) or XIAP deficiency, it is a rare form of primary immunodeficiency. Hallmarks of XLP-2 associated HLH are hemophagocytosis, cytopenia (pancytopenia, bicytopenia, thrombocytopenia or anemia), splenomegaly, fever and frequently chronic hemorrhagic colitis with cholangitis and/or hepatitis(Filipovich et al., 2010; Schmid et al., 2011). The clinical phenotype is associated with aberrant activation of macrophages and dendritic cells, and the accumulation of activated T lymphocytes in response to viral infections(Marsh et al., 2013). Cellular immunactivation is likely to be linked to the elevation of proinflammatory cytokines including IL-1β, IL-18, TNF, interferon-γ and IL-6 (Marsh et al., 2010; Rigaud et al., 2006; Wada et al., 2014).

Therefore, we sought to understand the role of direct TNFR2 activation in the absence of XIAP. Here, we provide evidence that specific TNFR2 stimulation primes macrophages for the upregulation of inflammasome components in both wildtype and *xiap*^−/−^ macrophages. Subsequent TNF release causes the activation of TNFR1 and then leads to gasdermin D mediated pyroptosis dependent on RIPK1 kinase activity and ROS production. While we were able to block cell death and therefore many of the cytokines returned to baseline, IL-18 levels did not. Thus, separating a key cytokine implicated in the etiology of XIAP deficient patients from cell death. Taken together, we discovered a novel role of XIAP to inhibit TNFR2 induced inflammatory pyroptosis and thereby suggest TNFR2 as a possible new therapeutic target in XLP-2 patients to dampen inflammation.

## Results

### Loss of XIAP RING domain sensitizes macrophages and neutrophils to TNFR2 induced cell death

We previously observed that only *xiap*^−/−^ bone marrow derived macrophages (BMDMs) were sensitized to cIAP1/2 specific Smac mimetics (Birinapant) induced cell death(Wong et al., 2014) and were surprised to find that the use of multimeric TNF (mega TNF) alone caused cell death (Figure 1A). To assess if specific activation of TNFR1 or TNFR2 was responsible, we used human TNF (TR1-TNF) to activate only TNFR1 and the recently published nonameric TNF fusion protein, TNC-sc(mu)TNF80 (TNC-TNF), to specifically activate TNFR2(Bossen et al., 2006; Rauert et al., 2010). We treated BMDMs from *xiap*^−/−^, *ciap1*^−/−^ and *ciap2*^−/−^ mice with either TR1-TNF or TNC-TNF overnight and assayed cell death by propidium iodide (PI) incorporation on flow cytometry (Figure 1B). Interestingly, *xiap*^−/−^ macrophages but not *ciap1*^−/−^ or *ciap2*^−/−^ macrophages were sensitive to TNFR2 induced cell death but insensitive to TNFR1 stimulation. The same was seen when we stimulated freshly isolated macrophages (CD11b^+^F4/80^+^) from *xiap*^−/−^ bone marrow (Figure S1A). To determine the kinetics of the observed cell death, BMDMs were imaged over the course of TNC-TNF treatment using time-lapse photography for the uptake of PI. *Xiap*^−/−^ macrophages started to die by 8h after TNC-TNF treatment as shown by the increase in PI positivity compared to wildtype cells (Figure 1C and S1B).

**Figure 1.**
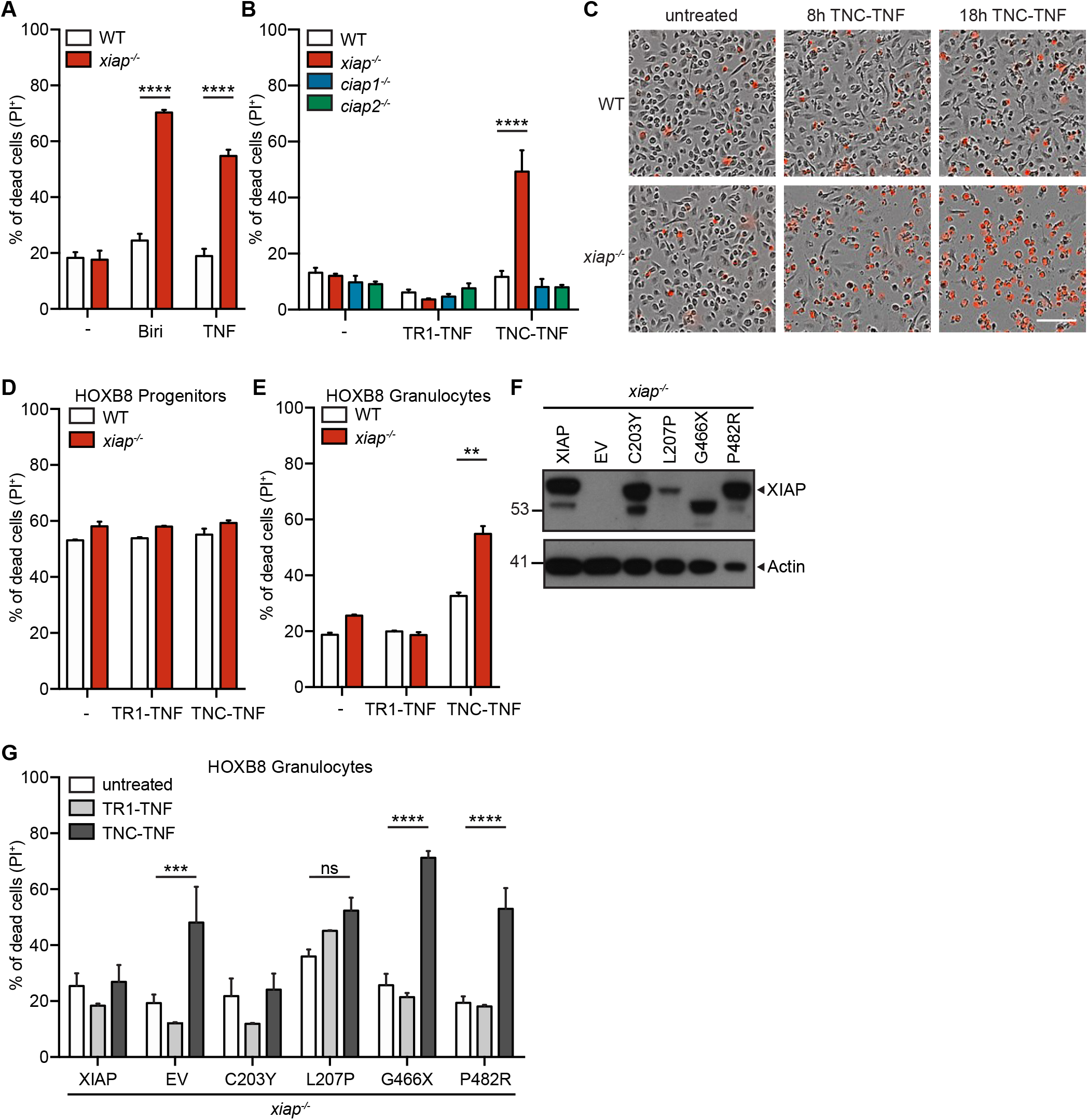
Loss of XIAP results in TNFR2 mediated cell death in myeloid cells. (A, B, D-F) Cell death was measured by the uptake of Propidium Iodide (PI) and analyzed by flow cytometry. (A) BMDMs from wildtype (WT) and *xiap*^−/−^ were stimulated with Birinapant (Biri) and multimeric TNF (human mega TNF) overnight. (B) BMDMs from WT, *xiap*^−/−^, *ciap1*^−/−^, and *ciap2*^−/−^ mice were treated overnight with TR1-TNF or TNC-TNF. (C) Representative phase contrast image merged with PI positive image at 0, 8 and 18 hours after TNC-TNF stimulation (3 independent experiments performed). (D) WT and *xiap*^−/−^ HOXB8 progenitors and (E) differentiated HOXB8 granulocytes were treated with TR1-TNF or TNC-TNF for 24h. (F) *Xiap*^−/−^ HOXB8 granulocytes were transfected with either WT, XIAP or XIAP mutated constructs and stimulated with TR1-TNF or TNC-TNF for 24h. (G) Basal expression levels of XIAP in HOXB8 progenitors were assessed by western blot. Blot is a representative of 3 independent experiments. Data shown is mean ± SEM including n=3-5 biological replicates. Experiments were repeated at least three times independently. Statistical significance was calculated using (A, B, D, E) two-way ANOVA with p **=<0.01, ****=<0.0001 or (F) a minimum of three independent experiments, including triplicates for each experiment. Statistical significance was calculated using Student-t test with p***=<0.001, ****=<0.0001.

There are various mutations found in all over the gene encoding XIAP, contributing to immune hyper-activation and tissue inflammation in XLP-2 patients(Prokop et al., 2017). To determine whether these mutations in XIAP cause sensitivity to TNFR2 induced cell death, we utilized the HoxB8 progenitor system(Wicki et al., 2016). Briefly, lineage negative cells from the bone marrow were transduced with 4-hydroxytamoxifen (4-OHT) inducible HoxB8 and immortalized with stem cell factors. Upon withdrawal of 4-OHT, progenitor cells differentiate into granulocytes. Progenitor cells were assayed for sensitivity to TNFR1 or TNFR2 specific stimulation. Both wildtype and *xiap*^−/−^ HoxB8 progenitor cells were insensitive to cell death (Figure 1D). However, differentiated *xiap*^−/−^ granulocytes were sensitive to TNFR2 induced cell death but not to TR1-TNF (Figure 1E). To assess the contribution of either the BIR or RING domain of XIAP to the observed cell death, different XIAP constructs containing mutations found in XLP-2 patients were re-introduced into the HoxB8 progenitor cells(Wicki et al., 2016) (Figure 1F). *Xiap*^−/−^ HoxB8 granulocytes with XIAP wildtype reintroduced (*xiap*^+/+^) remained insensitive to either TR1- or TNC-TNF induced cell death (Figure 1G). Intriguingly, C203Y and L207P mutations in the BIR2 domain reduced the sensitivity of HoxB8 granulocytes to TNC-TNF while G466X and P482R were unable to rescue TNFR2 induced cell death (Figure 1G). These data suggest that activation of TNFR2 in the absence of XIAP, specifically the E3 ligase domain, results in cell death in the myeloid compartment.

### TNFR2 mediated cell death in XIAP deficient macrophages is RIPK1 kinase activity dependent but independent of downstream necroptotic machinery or apoptosis

To determine the mode of cell death, we treated wildtype and *xiap*^−/−^ macrophages with TNC-TNF in combination with a pancaspase inhibitor (QVD) to assess the role of apoptosis or necroptosis with an MLKL inhibitor (GW806742X). TNFR2 induced cell death was slightly enhanced in the presence of the caspase inhibitor, suggesting the switch of the cell from apoptosis to necroptosis. Indeed, the MLKL inhibitor blocked TNFR2 induced cell death in *xiap*^−/−^ BMDMs (Figure 2A). In line with this, we did not detect any caspase activity upon TNC-TNF stimulation (Figure 2B) using a Smac mimetic (Compound A) that is known to induce apoptosis as control(Wong et al., 2014). To genetically confirm the role of necroptosis in TNFR2 induced cell death, *xiap*^−/−^ mice were crossed with *ripk3*^−/−^ or *mlkl*^−/−^ mice. Surprisingly, cell death analysis via time-lapse photography showed that in the absence of XIAP the loss of RIPK3 or MLKL did not reduce the TNC-TNF induced cell death (Figure 2C and Figure 2SA). In addition, at 8h the co-incubation with QVD did not rescue *xiap*^−/−^*ripk3*^−/−^ or *xiap*^−/−^*mlkl*^−/−^ from TNFR2 mediated cell death, suggesting no switch from apoptosis to necroptosis or vice versa when one pathway is blocked (Figure 2SB). These data suggest that neither caspase-mediated apoptosis nor RIPK3/MLKL-mediated necroptosis is triggered in response to TNFR2 specific activation in XIAP deficient macrophages. This is in contrast to LPS induced cell death where it has been shown to be RIPK3 and caspase-8 dependent in the absence of XIAP(Lawlor et al., 2017).

**Figure 2.**
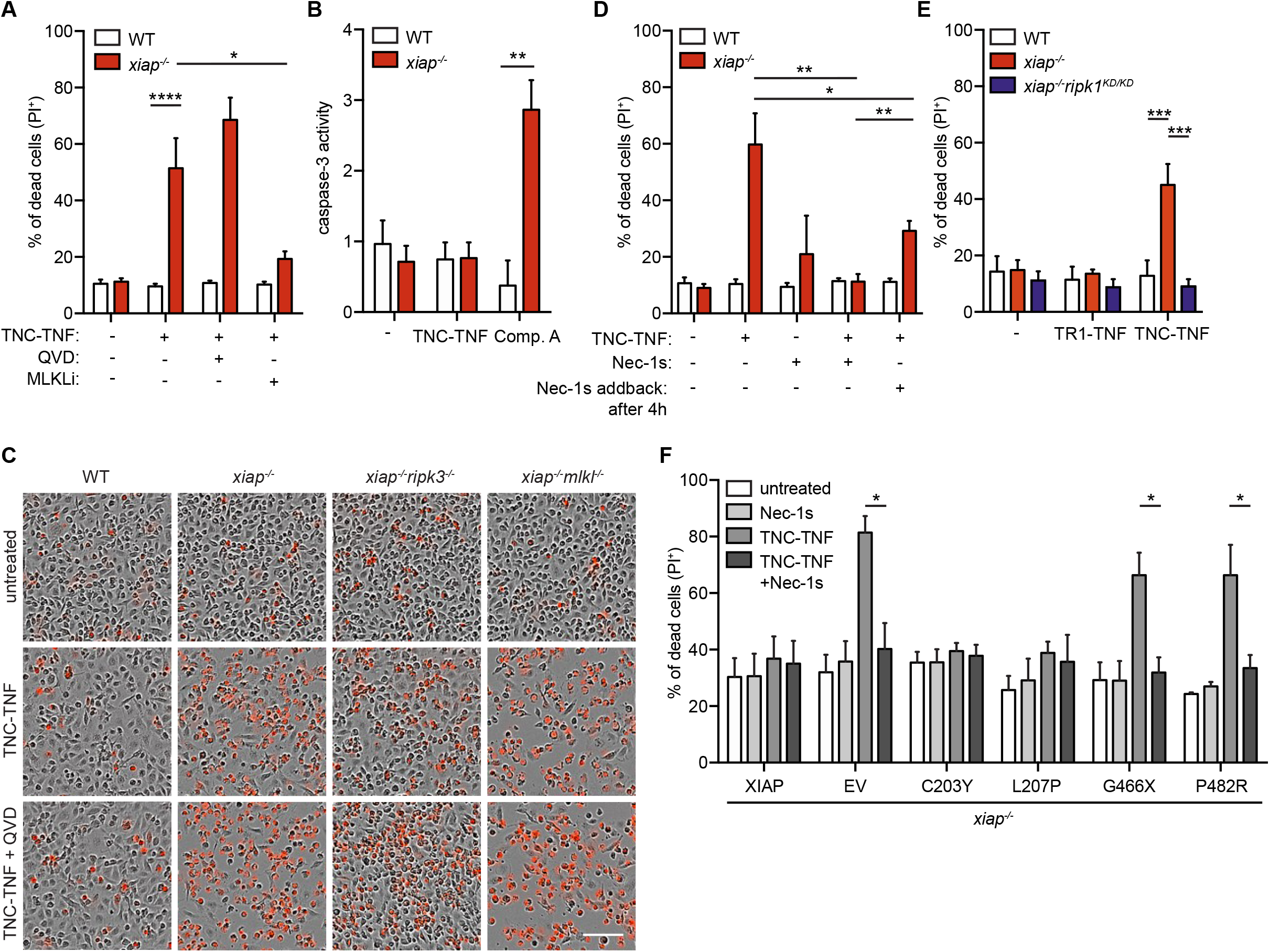
RIPK1 kinase activity mediates TNFR2 induced cell death in *xiap*^−/−^ macrophages, independent of apoptosis and necroptosis. (A) WT or *xiap*^−/−^ BMDMs were treated overnight with TNC-TNF in combination with either the caspase inhibitor QVD or MLKL inhibitor (GW806742X). Cell death was measured via PI uptake by flow cytometry. (B) BMDMs were treated with TNC-TNF or Compound A (positive control) for 12h, cells were lysed and caspase activity was measured over time using a fluorogenic DEVD AMC assay. Caspase activity is represented as fold change over untreated. (C) Representative phase contrast image at 18 hours after TNC-TNF and QVD stimulation in WT, *xiap*^−/−^, *xiap*^−/−^*ripk3*^−/−^, *xiap*^−/−^*mlkl*^−/−^ BMDMs monitored for cell death by PI uptake using time-lapse photography. (D) Cells were treated with TNC-TNF in combination with the RIPK1 kinase inhibitor Necrostatin-1s (Nec-1s) or TNC-TNF pre-stimulation for 4h and additional Nec-1s treatment. After 24h cell death was measured by PI uptake and analysed by flow cytometry. (E) WT, *xiap*^−/−^ or *xiap*^−/−^*ripk1^KD/KD^* BMDMs were treated with TNC-TNF and after 24h via PI uptake by flow cytometry. (F) *Xiap*^−/−^ HoxB8 granulocytes were transfected with either WT, XIAP or XIAP mutated constructs and stimulated with TNC-TNF or in combination with Nec-1s for 24h. Data shown is mean ± SEM of a minimum of three independent experiments, including triplicates for each experiment. Statistical significance was calculated using Student-t test with p *=<0.05. (A, B, D, E) Data shown is mean ± SEM including n=3-5 biological replicates. Experiments were repeated at least three times independently. Statistical significance was calculated using two-way ANOVA with p *=<0.05, **=<0.01, ***=<0.001, ****=<0.0001.

RIPK1 has been implicated in the regulation of cell death, via RIPK3 or its complex formation with caspase-8(Feoktistova et al., 2011; Tenev et al., 2011). Furthermore, it’s kinase activity has been shown to be important in the regulation of TNF production(Berger et al., 2014; Najjar et al., 2016; Wong et al., 2014). Intriguingly, even though we excluded RIPK3 and MLKL mediated necroptosis, when we stimulated wildtype and *xiap*^−/−^ macrophages with TNC-TNF we observed pronounced phosphorylation of RIPK1 at Ser166, which is indicative of necroptosis(Lin et al., 2016; Newton et al., 2016) (data not shown). Therefore, we were interested whether the kinase activity of RIPK1 is involved in TNFR2 induced cell death and stimulated wildtype and *xiap*^−/−^ macrophages with TNC-TNF in combination with a selective kinase inhibitor of RIPK1 (Necrostatin-1s, Nec-1s). The TNC-TNF induced cell death was entirely rescued when RIPK1 kinase activity was inhibited and even after 4h of pre-stimulation with TNC-TNF the inhibitor significantly rescued cell death (Figure 2D). To confirm our findings and exclude any possible off-target effect of Necropstatin-1s, we crossed *xiap*^−/−^ with a mouse strain containing a kinase dead mutant of RIPK1, *ripk1^K45A/K45A^* (*xiap*^−/−^*ripk1^KD/KD^*) (Berger et al., 2014). Consistent with Necrostatin-1s, macrophages from *xiap*^−/−^*ripk1^KD/KD^* mice were insensitive to TNC-TNF induced cell death (Figure 2E).

HoxB8 granulocytes were similarly resistant to TNC-TNF in the presence of Nec-1s as macrophages (Figure 2F). Taken together, TNFR2 induced cell death in the absence of XIAP is consistent between other myeloid cell types such as granulocytes and is dependent on RIPK1.

### TNFR2 stimulation leads to soluble TNF production resulting in TNFR1 mediated cell death in *xiap*^−/−^ macrophages

Our previous data and others suggested that the inhibition or loss of XIAP, cIAP1 and cIAP2 leads to TNF production which may lead to TNFR1 mediated cell death(Siegmund et al., 2016; Wong et al., 2014). To test if TNFR2 activation resulted in subsequent release of TNF and TNFR1 activation, we stimulated macrophages for 4h with TNC-TNF and subsequently added anti-TNFα to neutralize any TNF being produced upon the stimulation. This indeed reduced the amount of TNC-TNF induced cell death significantly, the reduction being ~30% (Figure 3A). We could further confirm the importance of autocrine TNF by stimulating *xiap*^−/−^ *tnf*^−/−^ macrophages with TNC-TNF, which were found to be resistant to TNC-TNF induced cell death (Figure 3B). To determine if soluble TNF was produced in response to TNC-TNF, we utilized *tnfr1*^−/−^*tnfr2*^−/−^ fibroblasts with re-introduced TNFR1. The cytoplasmic portion of TNFR1 was replaced with Fas and activation of TNFR1 results in cell death(Krippner-Heidenreich et al., 2002). Supernatant transfers from wildtype and *xiap*^−/−^ macrophages treated with TNC-TNF to TNFR1-Fas mouse fibroblast cells (TNFR1-Fas MFs) showed TNFR1 mediated cell death induced by soluble TNF which is produced by XIAP deficient macrophages (Figure S3A). To identify the importance of TNFR1 in the cell death seen, we differentiated BMDMs from *xiap*^−/−^*tnfr1*^−/−^ mice and found that they were insensitive to TNC-TNF induced cell death (Figure 3C). TNF mRNA production in response to TNC-TNF was independent of TNF/TNFR1 mediated feedback signaling as *xiap*^−/−^*tnfr1*^−/−^ macrophages produced similar levels of TNF mRNA in response to TNC-TNF compared to *xiap*^−/−^ BMDMs (Figure 3D). These results suggest that TNFR2 specific activation initiates a cell death via TNF/TNFR1 in the absence of XIAP.

**Figure 3.**
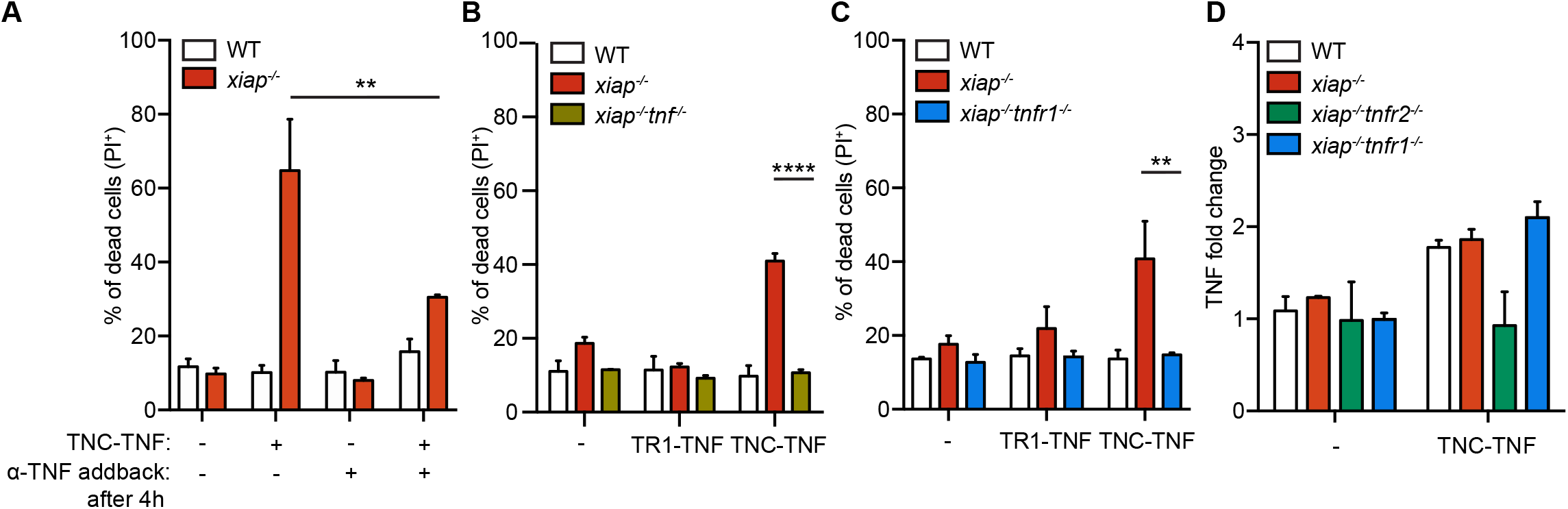
TNFR2 induced cell death in *xiap*^−/−^ macrophages requires soluble TNF and TNFR1 activation. (A) BMDMs from WT and *xiap*^−/−^ were treated with TNC-TNF and/or anti-TNFα after 4h of stimulation and after 24h, cell death was measured via PI uptake by flow cytometry. (B,C) *xiap*^−/−^*tnf*^−/−^ and *xiap*^−/−^*tnfr1*^−/−^ BMDMs are resistant to TNC-TNF induced cell death.(D) Relative TNFα mRNA levels were compared after 2h of TNC-TNF or TNC-TNF and Nec-1s treatment from BMDMs of the indicated genotypes. Data shown is mean ± SEM including n=3-5 biological replicates. Experiments were repeated at least three times independently. Statistical significance was calculated using two-way ANOVA with p *=<0.05, **=<0.01, ***=<0.001, ****=<0.0001.

### TAK1/RIPK1 mediated TNF production downstream of TNFR2 activation

Subtle changes in TNFR1 signaling in the absence of XIAP have been identified such as delayed(Yabal et al., 2014). To determine if XIAP influenced downstream signaling of TNFR2, we probed for changes in NF-κB and MAPK pathways. In *xiap*^−/−^ macrophages, we found a diminished phosphorylation of ERK (p-ERK) and prolonged phosphorylation of p65 (p-p65) representing either MAPKs or NF-κB signaling pathways, respectively (Figure 4A). Phosphorylation of p38 and JNK was not altered. These data suggest that the direct activation of TNFR2 leads to changes in transcription in the absence of XIAP. To determine the significance of the changes in NF-κB, we used IKK inhibitor VII and TAK1 inhibitor (5Z-7-Oxozeaenol) and MAPK pathways (SCIO469, p38 inhibitor; PF3644022, ERK1/2 inhibitor) and co-incubated with TNC-TNF. TNF mRNA was assessed after 2 hours of stimulation. Surprisingly, only TAK1 inhibition led to a decrease of TNF production in response to TNFR2 (Figure 4B). Because our previous work showed that RIPK1 kinase inhibition could reduce TNF production in response to Smac mimetics(Wong et al., 2014), we next explored whether RIPK1 was involved in the production of TNF. Interestingly, the loss of RIPK1 kinase activity resulted in a reduction of TNF production but only in the absence of XIAP (Figure 4C). These results show that TAK1 as well as RIPK1 are downstream of TNFR2 similar to TNFR1 activation.

**Figure 4.**
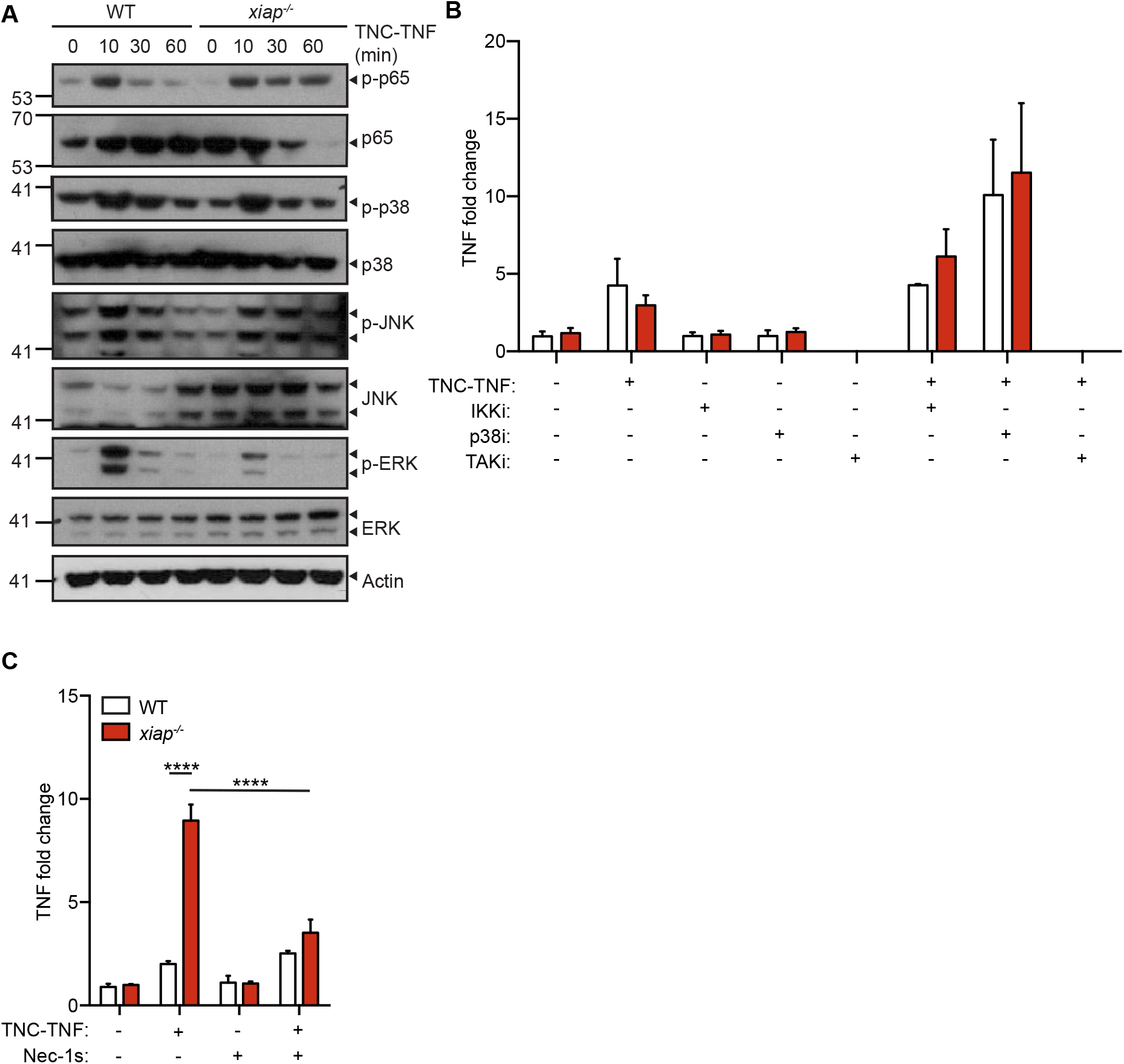
TNC-TNF prolongs signaling activation in *xiap*^−/−^ macrophages that is dependent on RIPK1 kinase activity. (A) BMDMs were treated with TNC-TNF for indicated time points and expression levels of proteins of the NF-κB and MAPK signaling pathway were analysed by western blotting. (B) Relative TNFα mRNA levels were compared after 2h of TNC-TNF and/or IKK inhibitor, p38 inhibitor and TAK inhibitor treatment from WT and *xiap*^−/−^ BMDMs. (C) Relative TNFα mRNA levels were compared after 2h of TNC-TNF or TNC-TNF and Nec-1s treatment from BMDMs of the indicated genotypes. Data shown is mean ± SEM including n=3-5 biological replicates. Experiments were repeated at least three times independently. Statistical significance was calculated using two-way ANOVA with p ****=<0.0001.

### Response to TNFR2 shows similar enrichment of target genes compared to TNF/TNFR1 induction

To examine XIAP specific transcriptional changes, we performed RNA sequencing of WT, *xiap*^−/−^ and *xiap*^−/−^*tnfr1*^−/−^ macrophages treated with TNC-TNF for 2h. Comparison of the untreated samples of each genotype showed the gene expression was surprisingly similar (Figure 5A and Supp Table 1). Upon TNC-TNF stimulation, we found a set size of 258 genes in WT, 296 in *xiap*^−/−^ and 223 genes in *xiap*^−/−^*tnfr1*^−/−^ BMDMs that were differentially regulated (false discovery rate ≤ 0.05). The majority of genes were up-regulated in all three genotypes stimulated with TNC-TNF. Interestingly, there were 88 genes uniquely regulated by the loss of XIAP, 47 in wildtype and 32 genes in *xiap*^−/−^*tnfr1*^−/−^ macrophages. Only 15 genes were differentially regulated and overlapping between *xiap*^−/−^ and *xiap*^−/−^*tnfr1*^−/−^ macrophages when treated with TNC-TNF, suggesting changes in this set of genes is not influenced by TNFR1.

**Figure 5.**
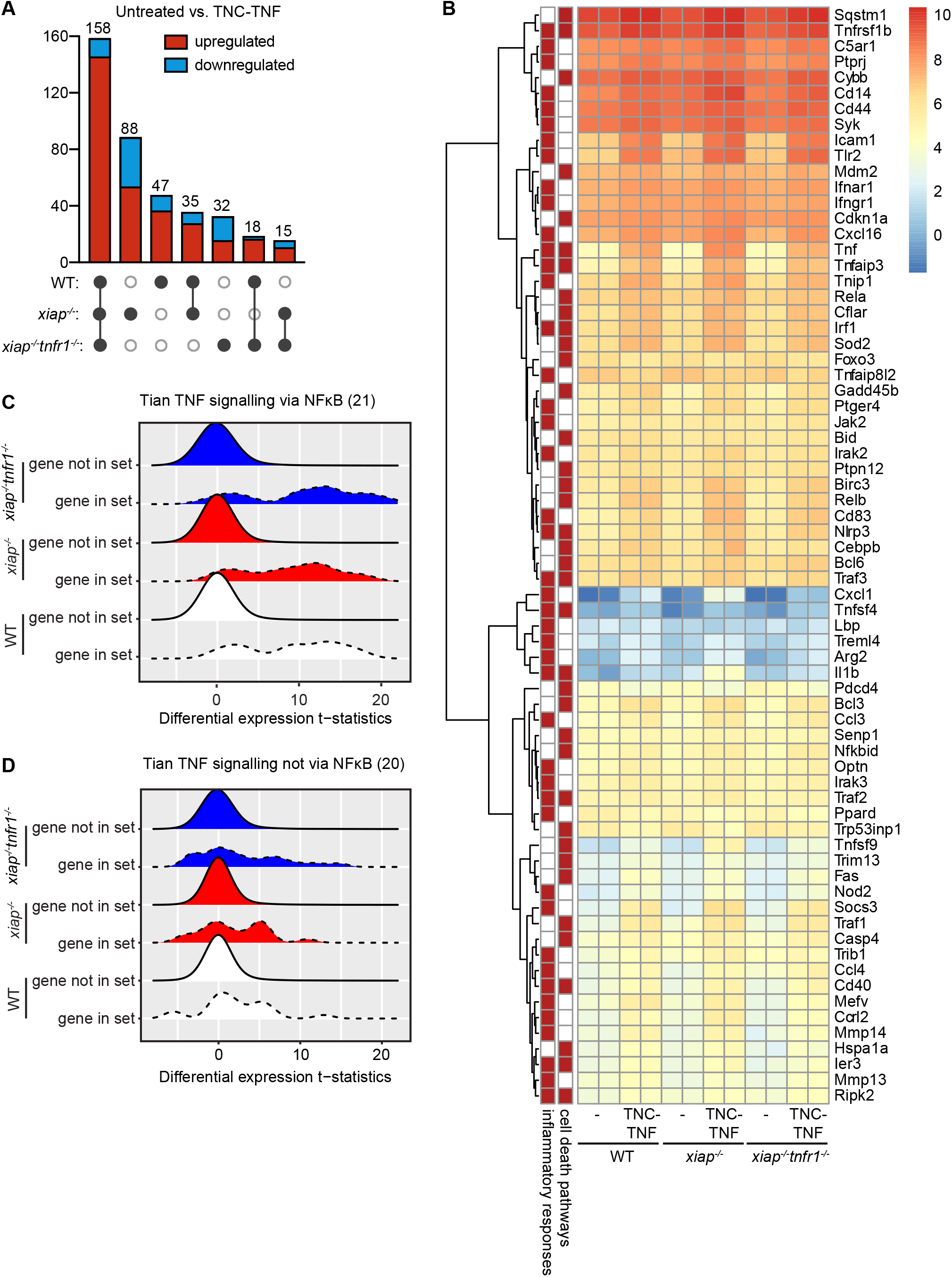
RNASeq analsysis of TNFR2-mediated gene signature in wildtype, *xiap*^−/−^ and *xiap*^−/−^*tnfr1*^−/−^ macrophages. BMDMs from WT, xiap−/− and xiap−/−tnfr1−/− were treated for 2h with TNC-TNF and transcriptional profiling of the RNA was performed. Genes that were differentially regulated with FDR >0.05 were enriched. (A) Upset plot representing the total differential regulation of genes compared between the genotypes, including up- and downregulated genes. (B) Differential regulated genes were analyzed for biological function including inflammatory responses and cell death using gene ontology and represented by hierarchical clustering in a heatmap. (C) Tian pathway analysis of genes significantly altered in all genotypes compared to TNF signaling via NF-κB or (D) not via NF-κB.

Gene ontology analysis showed an enrichment of genes involved in inflammation and cell death processes. These were clustered for expression based on genotype and interestingly, a similar pattern of up-regulation independent of genotype was seen (Figure 5B). Gene set enrichment analysis showed that the differentially regulated genes upon TNFR2 stimulation are alike to those associated with TNF/TNFR1 and NF-kB and to a lesser extent TNF signaling that is not reliant on NF-kB activation (Figure 5C and Supp Table 2). Genes up-regulated or down-regulated in LPS gene signatures, KEGG NOD-like receptor signaling and inflammatory signature were also significantly enriched in our gene sets showing the cross over of pathways linking LPS and TNF induction (Supp Table 2). These data surprisingly show that TNFR2-induced transcriptional changes in the absence of XIAP was minimal but that TNFR2 itself is a driver for inflammation in macrophages.

### Pro-inflammatory cytokines instigated by TNFR2 are dependent on TNF and RIPK1 kinase activity

Because TNFR2 is not normally associated with inflammation, we screened for cytokines/chemokines. From this assay, we found IL-10, IL-1β, IL-6, IL-18 and chemokines, CCL3, CCL4, CXCL1 and CXCL2 were further up-regulated in *xiap*^−/−^ macrophages compared to wildtype when treated with TNC-TNF for 24h (Figure S4A). CCL2, CCL5 and CCL7 were up-regulated to similar levels in both wildtype and *xiap*^−/−^ macrophages in response to TNF-TNF (data not shown). Because IL-1β, IL-18 and IL-6 have been identified in serum of patients during a HLH episode, we asked whether these cytokines were regulated in response to TNF. Using *xiap*^−/−^*tnf*^−/−^ BMDMs, we assayed for these cytokines after 12h of TNFR2 stimulation. IL-1β, IL-18 and Il-6 were dependent on TNF production (Figure 6A).

**Figure 6.**
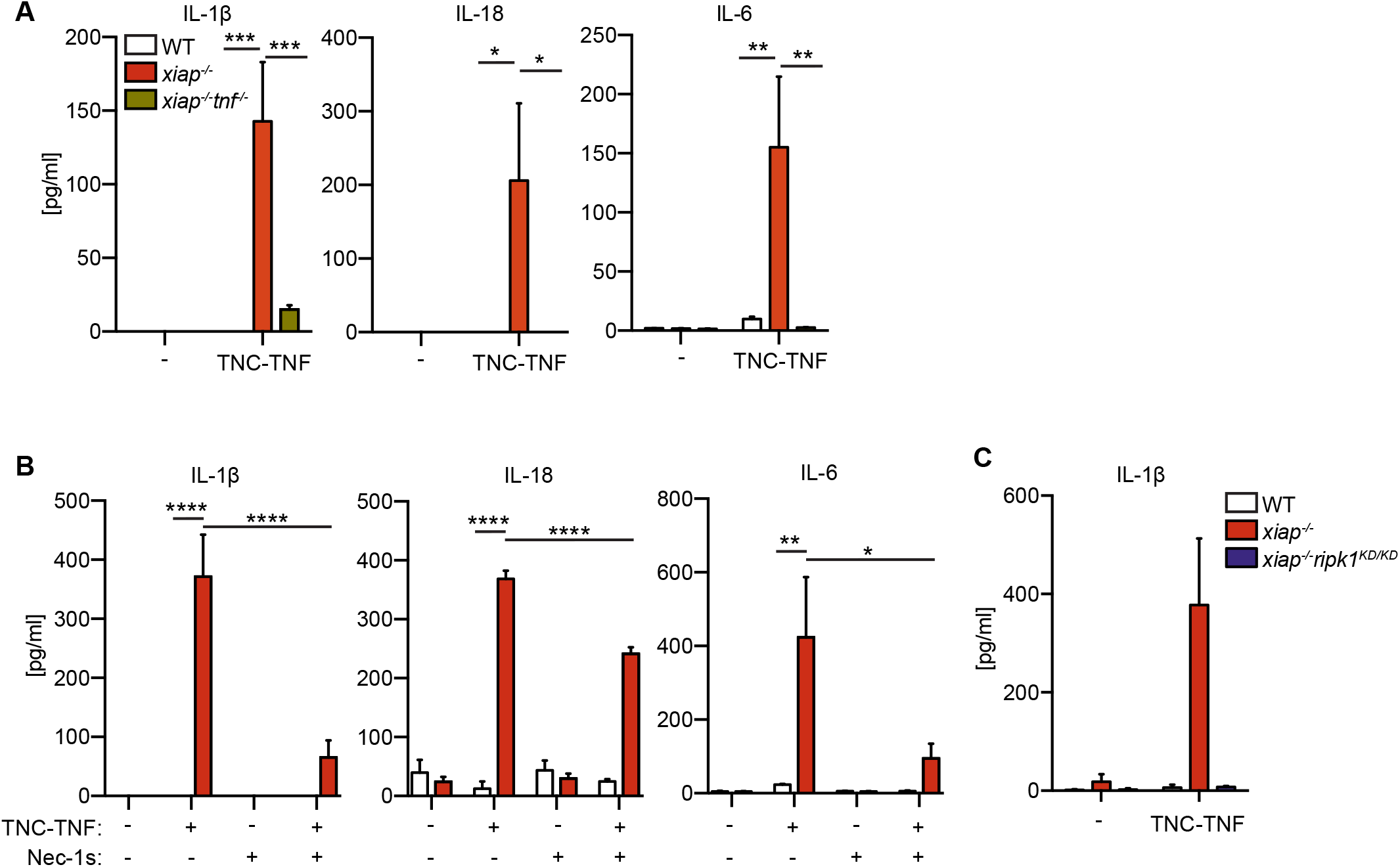
RIPK1 kinase activity regulates cytokine and chemokine production in *xiap*^−/−^ macrophages. (A) WT, *xiap*^−/−^, *xiap*^−/−^*tnf*^−/−^ BMDMs were stimulated with TNC-TNF and supernatants were assayed for cytokines by multiplex. After 12h supernatant was taken and assayed for IL-1β, IL-18 and IL-6. (B) BMDMs were treated overnight with TNC-TNF and/or Nec-1s and supernatant was assayed for IL-1β, IL-18 and IL-6. (C) BMDMs of WT, *xiap*^−/−^ and *xiap*^−/−^*ripk1^KD/KD^* treated with TNC-TNF for 12h and showed reduced IL-1β production upon loss of RIPK1 kinase activity. Data shown is mean ± SEM including n=2-4 biological replicates. Experiments were repeated at least three times independently. Statistical significance was calculated using two-way ANOVA with p *=<0.05, **=<0.01, ***=<0.001, ****=<0.0001.

We then asked whether inhibition of the RIPK1 kinase activity by Nec-1s also reduced the secretion of pro-inflammatory cytokines at 24h post stimulation with TNC-TNF. The reduction of IL-1β and IL-6 was reduced and not significant compared to basal levels while IL-18 levels remained significantly above basal levels (Figure 6B). Chemokine, CXCL1 was similarly down-regulated by co-incubation of Nec-1s with TNC-TNF (Figure S4B). IL-1β production was entirely diminished when treating *xiap*^−/−^*ripk1^KD/KD^* BMDMs, correlating with the Nec-1s data (Figure 6C). RIPK3 has been shown to limit IL-1β in the absence of XIAP in response to LPS(Vince et al., 2012; Yabal et al., 2014). Likewise, we found IL-1β but not IL-6 was reduced in *xiap*^−/−^*ripk3*^−/−^ and *xiap*^−/−^*mlkl*^−/−^ macrophages in response to TNC-TNF for 24h (Figure S4C). These results imply direct TNFR2 activation in the absence of XIAP leads to increased pro-inflammatory IL-1β, IL-18 and IL-6 in a RIPK1 kinase dependent manner.

### XIAP restricts the activation of TNFR2 induced priming of the pyroptotic machinery

We examined the gene expression of known cytokines, chemokines, and receptors previously implicated in cell death of XIAP deficiency or TNFR2 induced cell death. Consistent with other groups(Siegmund et al., 2018), we also found, *tnfaip3* (A20) and *traf1* up-regulated by TNC-TNF in our RNAseq analysis (Supp Table 1). The anti-apoptotic response of *cflar* (c-FLIP) and *birc3* (cIAP2) is consistent in *xiap*^−/−^ and *xiap*^−/−^*tnfr1*^−/−^ macrophages (Figure 7A). The log fold change plots show that while *tnf, cxcl1, cxcl2 and ccl3* are differentially regulated at the RNA level in all genotypes, there is a slight increase in RNA levels in the absence of XIAP (Figure 7A, coded in black). *Ccl4* and *il-1β* were exclusively up-regulated in *xiap*^−/−^ macrophages (Figure 7A, coded in grey). These data suggest that while TNFR2 directly stimulates the expression of key cytokines and chemokines, the cytokine network is further enhanced by the absence of XIAP and the stimulation of TNFR1.

**Figure 7.**
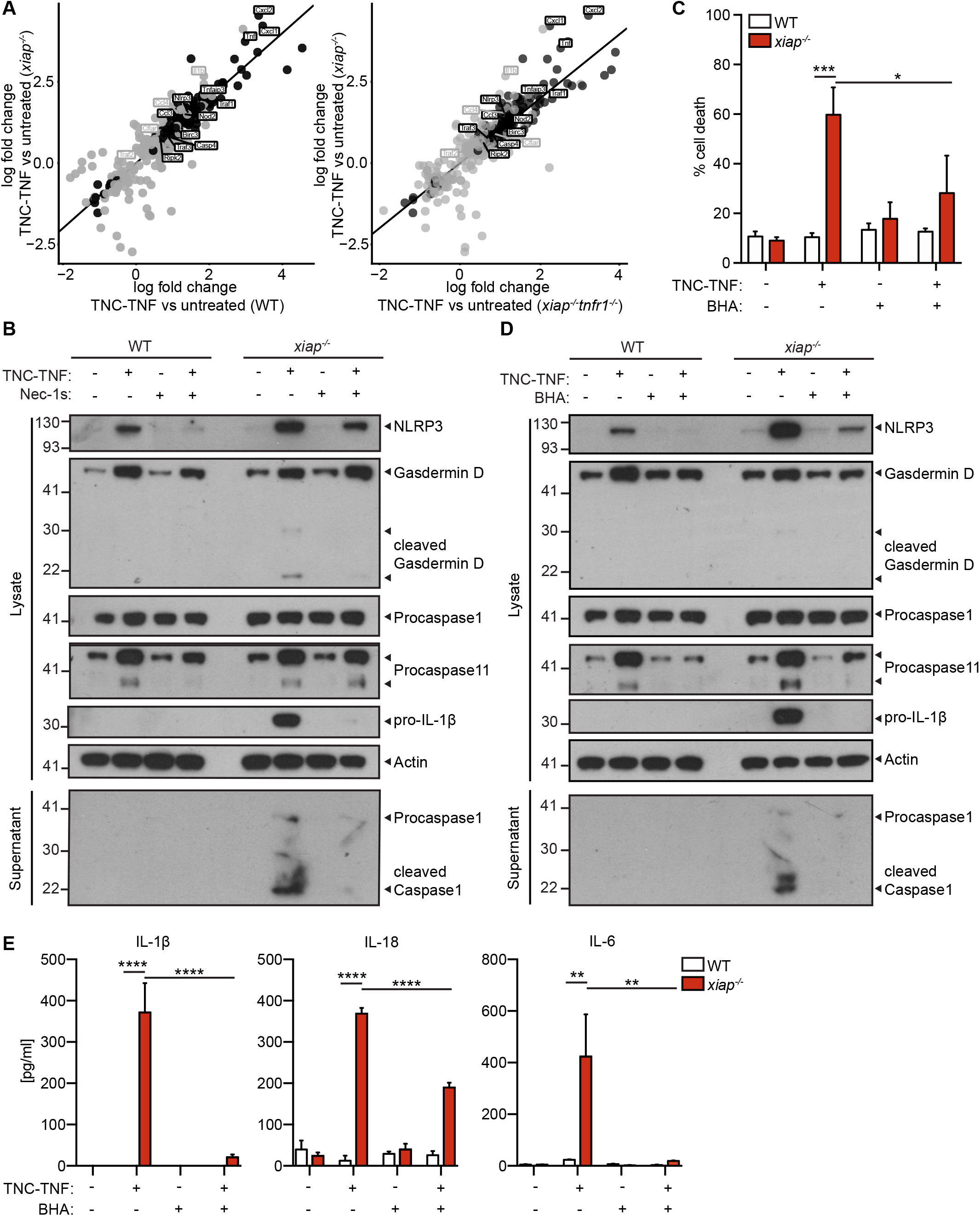
TNFR2 primes inflammasome components, NLRP3 and caspase-11, and in *xiap*^−/−^ macrophages, cleaved gasdermin D and increased IL-1β, leads to pyroptosis inhibited by RIPK1 kinase activity and BHA. (A) BMDMs from WT, *xiap*^−/−^ and *xiap*^−/−^*tnfr1*^−/−^ were treated for 2h with TNC-TNF and transcriptional profiling of the RNA was performed. Genes that were differentially regulated with FDR <0.05 were enriched. Log fold plot comparison of significantly differential genes between genotypes. Black represents the differential expression common in all genotypes and gray exclusively between the two genotypes depicted in the graph. (B) BMDMs from WT and *xiap*^−/−^ were treated overnight with TNC-TNF and/or BHA. Cell death was assessed by PI uptake on flowcytometry. (C) Cell lysates and supernatant of macrophages treated with TNC-TNF and Nec-1s or (D) BHA (5μM) were immunoblotted and probed for gasdermin D, NLRP3, caspase 11, caspase-1 and IL-1β. Western blots are a representative of at least 3 independent experiments. (E) BMDMs treated overnight with TNC-TNF in combination with BHA (5μM). IL-1β, IL-18 and IL-6 were measured in the supernatant of by multiplex. Data shown is mean ± SEM, including n=3 biological replicates. Experiment was repeated three times independently. Statistical significance was calculated using two-way ANOVA with p**=<0.01, ***=<0.001, ****=<0.0001.

In our analysis we identified the up-regulation of inflammasome related components such as NLRP3 and caspase-11 (*casp4*). To confirm the involvement of the pyroptotic components in TNFR2 induced cell death, we treated WT and *xiap*^−/−^ BMDMs with TNC-TNF overnight and assayed the supernatant as well as the lysate. We found that indeed, upon TNC-TNF, NLRP3 and caspase-11 were up-regulated as well as gasdermin D (Figure 7B). However, the activation of caspase-1 and cleavage of gasdermin D was only observed in XIAP deficient cells. Interestingly, the loss of RIPK1 kinase activity resulted in a slight decrease in gasdermin D and caspase-11 protein levels as well as cleavage of gasdermin D. Taken together, TNFR2 primes macrophages for pyroptotic cell death by increasing RNA and protein levels of NLRP3, caspase-11 and gasdermin D.

We then asked how NLRP3 may be activated in response to TNFR2 stimulation in XIAP deficient cells. Because ROS production has been detected before in conjunction with NLRP3 activation, we examined whether a free radical scavenger such as BHA or NAC could prevent cell death. Co-incubation of TNC-TNF and BHA or NAC reduced cell death in the *xiap*^−/−^ macrophages (Figure 7C). Protein levels of NLRP3, gasdermin D and caspase-11 were also reduced and subsequent caspase-1 cleavage was missing (Figure 7D). The loss of caspase-1 activation resulted in significantly lower levels of IL-1β, IL-18 and IL-6 in the supernatant of treated *xiap*^−/−^ macrophages (Figure 7E). However, unlike IL-1β and IL-6, IL-18 levels remained higher than baseline. Taken together, in the absence of XIAP, the increased levels of the inflammasome components are activated and result in inflammatory cell death.

## Discussion

XIAP deficient patients are characterized with multi-organ inflammation triggered by viral infections. XLP-2 is associated with activated macrophages and lymphocytes and overexpression of pro-inflammatory cytokines including TNF(Wada et al., 2014). The current model proposes XIAP deficiency pre-disposes cells to either increased production of pro-inflammatory cytokines and/or cell death. In our study, we identified that specific activation of TNFR2 induces inflammation in XIAP deficient macrophages, these cytokines are similar to those identified in the serum of XLP-2 patients. TNFR2 activation specifically increased the expression of inflammasome components and in the absence of XIAP, gasdermin D mediated pyroptotic cell death regulated in a RIPK1 kinase activity and ROS dependent manner. While cell death could be blocked by RIPK1 kinase activity or attenuation of ROS production, only IL-18 was not reduced upon inhibition of cell death. Thus, our study provides evidence that inhibition of cell death will not result in reduction of key inflammasome cytokines, IL-18.

The concept of non-death domain TNF super family receptors (TNFSFR) involved in cell death has been previously reported in cancer cell lines for FN14, CD40, TNFR2, and CD30, acting through a TNF/TNFR1 axis(Grell et al., 1999; Schneider et al., 1999; Vince et al., 2008). Our results show that primary macrophages can undergo non-death domain TNFSFR induced cell death involving soluble TNF and TNFR1 but only in the absence of XIAP. Caspase inhibition slightly enhanced TNFR2-mediated cell death but did not shift the cell death to necroptosis involving RIPK3 or MLKL as reported previously by Siegmund *et al*. (2018) implying that caspase-8 activity is not required for cell death. Interestingly, our transcriptional profiling revealed that anti-apoptotic genes such as *cflar* (c-FLIP) and *birc3* (cIAP2) were up-regulated in *xiap*^−/−^ and *xiap*^−/−^*tnfr1*^−/−^ macrophages. Thus, it is possible that the increase in c-FLIP may stabilize complex II downstream of TNFR1, causing increased NF-κB activation. In this way, using a pan-caspase inhibitor in combination with TNFR2 stimulation may also stabilize complex II, thereby increasing and activating cell death (Siegmund et al., 2018; 2016).

Cell death induced by TNFR2 in *xiap*^−/−^ macrophages was only reduced by inhibition of RIPK1 kinase activity. We also observed the ability of RIPK1 to remain phosphorylated at Ser166, a known indicator for the activation of necroptosis(Ofengeim et al., 2015), was limited by XIAP. Yet, we found the inhibition of the kinase activity reduced TNFR2 mediated TNF mRNA production, which is needed to activate TNFR1 to mediate cell death. These data would suggest RIPK1 mediates inflammation, specifically TNF, in line with previous results linking the kinase activity of RIPK1 to inflammation(Berger et al., 2014; Najjar et al., 2016; Wong et al., 2014). Our addback studies however, do not exclude a role for RIPK1 kinase activity downstream of TNFR1 activation. This is in contrast with recent studies showing that LPS induced cell death and inflammasome activation in the absence of XIAP is dependent on RIPK3 but not RIPK1 kinase activity(Vince et al., 2012; Yabal et al., 2014).

While XIAP influences in TNF dependent cytokine production and cell death downstream of TNFR1(Yabal et al., 2014), alterations in NF-κB and MAPK signaling downstream of TNFR1 have not been reported. Here, we show that the signaling downstream of TNFR2 was altered with sustained p-p65 activation and decreased phosphorylation of ERK in *xiap*^−/−^ macrophages (Figure 6B). This suggests that XIAP acts downstream of TNFR2 rather than TNFR1, modulating MAPK and NF-κB activation. XIAP has also been shown to bind TAB1 through its BIR1 domain and thereby controlling TAK1 and subsequent NF-κB activation(Lu Mol Cell 2007; Hofer-Warbinek et al., 2000). We found that the RING function of XIAP mediates TNFR2 induced cell death. These results differ from MDP/NOD2 mediated NF-κB activation, which has been linked to both mutations in the BIR2 and RING domain of XIAP(Chirieleison et al., 2017; Damgaard et al., 2013; 2012). Further studies are required to understand how XIAP is interacting with signaling complexes downstream of TNFR2.

Despite the changes in signaling observed downstream of TNFR2 activation, the transcriptional profile response was surprisingly similar between wildtype, *xiap*^−/−^ and *xiap*^−/−^ *tnfr1*^−/−^ macrophages. The genes commonly altered in the different genotypes were enriched for inflammation, immune and cell death. We detected an increase in eight different cytokines and chemokines in *xiap*^−/−^ macrophages treated with TNFR2 (CCL2, CCL3, CCL4, CXCL1, CXCL2, Eotaxin, IL-1β, IL-6, IL-18) and some but not all were also found up-regulated in the RNAseq data (*ccl2, ccl3, cxcl1, cxcl2* and *il-1β)*. The transcriptional profiling revealed that in the absence of TNFR1, despite no cell death, many of these cytokine and chemokines were similarly up-regulated compared to wildtype. This indicates after TNFR2 activation, that subsequent stimulation of TNFR1 signaling may aid in additional increase at the RNA and protein level. The production of TNF however is direct from TNFR2 stimulation as the upregulation of TNF at the RNA level was evident in xiap−/−tnfr1−/− macrophages. Alternatively, the presence of cell death enhanced the cytokine production through DAMPs. More importantly, the increase in IL-1β, IL-18 and IL-6 only in the absence of XIAP, resembles those identified in the serum of XIAP deficient patients implying TNFR2 activation is a critical part in the initiation of the cytokine storm.

The RNAseq data further revealed TNFR2 induced inflammasome components, NLRP3 and caspase-11. Protein levels were induced irrespective of genotype and likewise, RIPK1 kinase activity and ROS inhibition reduced NLRP3 and caspase-11. The increase in NLRP3 and caspase-11 protein expression is clearly independent of cell death induction as it happens in wildtype cells. Our data contrasts from the use of LPS to induce necroptosis/inflammasome activation in XIAP deficient cells. We did not find TNFR2-induced pyroptosis to be RIPK3 dependent(Lawlor et al., 2017; 2015; Yabal et al., 2014). The difference of those findings may lie in multiple pathways activated by LPS while we directly stimulated TNFR2 only. TNFR2 is not a traditional inflammasome stimulus, therefore the priming to activate the inflammasome may be the ROS production in the absence of XIAP or the addition of DAMPs due to the presence of cell death. Loss of RIPK1 kinase activity or ROS were able to limit inflammasome activation, cell death and the secretion of Il-1β but interestingly, IL-18 was far from baseline.

TNFR2 has been shown to promote survival, differentiation and promote a immune suppressive function (Polz et al., 2014; Zhao et al., 2012). However, our data suggest the opposite in the absence of XIAP. Here, we provide evidence that the activation of TNFR2 primes macrophages towards inflammasome activation with dominant expression of proinflammatory cytokines. Our findings are of therapeutic interest as XLP-2 patients have increased inflammasome related cytokines. Of the cytokines identified upon HLH episodes in XLP-2/XIAP deficient patients, IL-18 is one of the pro-inflammatory cytokines which does not reduce to baseline in the serum(Wada et al., 2014). Our study reveals a novel mechanism that in the complete absence of XIAP or a XIAP RING mutation, myeloid cells are primed for inflammasome activation through TNFR2 stimulation. RIPK1 kinase activity is required for soluble TNF production, TNFR1 and ROS production culminating in gasdermin D mediated pyroptosis. We hypothesize that infections stimulate TNFR2 to prime the inflammasome machinery and in the absence of XIAP convert the inflammation to HLH. Thus, we think direct targeting of TNFR2 but not TNFR1 can be of potential therapeutic interest in XIAP deficient patients to limit inflammation.

## Acknowledgements

This research was supported by Hartmann Müller Stiftung, ZUNIV-FAN, Swiss National Science Project Grant, SPARKS XLP Trust Fund, Stiftung für wissenschaftliche Forschung an der Universität Zürich, EMDO Stiftung, Krebsliga Zürich. LMS and LV were supported by the Forschungskredit Candoc (University of Zurich). WWW is supported by the Clöetta Medical Research Fellow. HW is supported by DFG (grants WA1025/31-1 and TRR221 project B02. PJJ was supported by a Max Eder-Program grant from the Deutsche Krebshilfe (program #111738), Deutsche José Carreras Leukämie-Stiftung (DJCLS R 12/22 and DJCLS 21R/2016), Else Kröner Fresenius Stiftung (2014_A185), from the Deutsche Forschungsgemeinschaft (DFG FOR 2036) and the Deutsche Konsortium für translationale Krebsforschung (DKTK) of the German Cancer Center (DKFZ). The authors thank Prof. Dr. C. Münz and A. Fontana for critical reading and discussions.

## Author contributions

JK, LMS and WWW designed the research. JK, LMS, SR, RR, LV and WWW performed the experiments. MY, PJJ, HW, MR and TK provided reagents and cell lines that made this study possible. JK, LMS, SR, MR and WWW analyzed the data. LMS and WWW wrote the manuscript. JK, SR, LV, MY, PJJ, HW, MR and TK gave feedback on the draft paper.

## Materials & Methods

### Mice

*Xiap*^−/−^, *ciap1*^−/−^, *ciap2*^−/−^ and *ciap1^LC^ciap2*^−/−^ mice were a kind gift from J. Silke from WEHI and were previously described(Moulin et al., 2012). These strains were embryo transferred and maintained in an SPF facility in Zurich. *Tnfr1*^−/−^ and *tnfr2*^−/−^ mice were a kind gift from A. Fontana and A. Aguzzi, respectively(Peschon et al., 1998) and were crossed to *xiap*^−/−^ mice to generate *xiap*^−/−^*tnfr1*^−/−^ and *xiap*^−/−^*tnfr2*^−/−^ mice. *Ripk1*^K45A/K45A^ (*ripk1^KD/KD^*) mice were a gift from GlaxoSmithKline and were crossed to *xiap*^−/−^ mice to generate *xiap*^−/−^*ripk1^KD/KD^* mice. All mice used in this study were back-crossed to C57BL/6 mice. All animal experiments were performed at the University of Zurich under the ethical license 186/2015. *Xiap*^−/−^*tnf*^−/−^, *xiap*^−/−^ *ripk3*^−/−^, *xiap*^−/−^*mlkl*^−/−^ mice were a kind gift from P. Jost and housed at the Technical University of Munich. Experiments were conducted in accordance with GSK policies on the care, welfare, and treatment of laboratory animals.

### Generation of bone marrow derived macrophages and cell lines

To generate bone marrow-derived macrophages (BMDMs), bone marrow was obtained from the tibia and femur of 6-12 weeks old mice. Cells were cultured on petri dishes for 5 days in Dulbecco’s modified Eagle medium (DMEM, Gibco) containing 1g/L glucose, 1% (v/v) penicillin/streptomycin/glutamine (Gibco), 10% FBS (SeraGlobe) and supplemented with 20% (v/v) L929 mouse fibroblast conditioned medium. On day 5, cells were harvested and seeded at 1×10^6^ cells/mL into the desired tissue culture plates (e.g. 1×10^5^ cells per 96 well). TNFR1-Fas and TNFR2-Fas expressing mouse fibroblasts were obtained from Anja Krippner-Heidenreich(Krippner-Heidenreich et al., 2002) and were cultured in DMEM containing 1g/L glucose, 1% (v/v) penicillin/streptomycin/glutamine and 10% FBS. HOXB8 progenitor cells were cultured in RPMI 1640 media with 10% (v/v) heat-inactivated FBS, 1% (v/v) penicillin/streptavidin, 7% (v/v) SCF from CHO/SCF conditioned medium and 1μM 4-hydroxytamoxifen (4-OHT, MedChemExpress)(Wicki et al., 2016). To differentiate granulocytes from the HOXB8 progenitors, cells were washed twice with PBS and resuspended at a concentration of 2.5×10^4^ cells/mL in media without 4-OHT and differentiated for 5 days.

### Ligands and inhibitors

To produce the agonistic TNFR2 ligands, pCR3 Fc-Flag-TNFR2-specific nonameric murine TNF variant (TNC-TNF), pCR3 Flag-TNC-TNF or pCR3 Fc-Flag human TNFR1 (TNFR1-TNF), were transfected into 293t cells and purified as previous described(Fick et al., 2012). Endotoxin levels were tested and removed (Pierce High Capacity Endotoxin Removal Spin columns, ThermoScientific). TNF variants were used at 100ng/mL. Inhibitors were used at the following concentrations: Birinapant (500nM, Chemietek), Compound A (500nM, Tetralogics), multimeric TNF (mega TNF, 100ng/mL, Adipogen) Necrostatin-1s (1μM, MedChemExpress), Q-VD-OPH (5μM, Adipogen), MLKLi (0.1μM, GW806742X, SYNkinase), SCIO469 (Tocris), IKKVII (Calbiochem), 5Z-7-Oxzeaenol (Sigma).

### Antibodies

The following antibodies were used for flow cytometry: CD11b-PeCy7 (clone M1/70, eBioscience), F4/80 (clone BM8, eBioscience), and fixable viability dye (eBioscience). The following antibodies were used for western blotting: phospho-p65, NF-κB2, phospho-ERK, phospho-JNK, phospho-p38, total ERK, total JNK and total p38 from Cell Signaling Technologies. Phospho-RIPK1 was received from Genentech. Total p65 was purchased from Santa Cruz. TRAF2, gasdermin D and caspase-11 were purchased from Abcam, caspase-1 from Adipogen and IL-1β from RnD. cIAP1 was purchased from Human Atlas. RIPK1 and XIAP was purchased from BD Biosciences. Secondary antibodies for western blotting such as donkey anti mouse/rabbit/rat IgG conjugated to HRP are from SouthernBiotec, the donkey anti goat IgG was purchased from Santa Cruz. The neutralizing antibody against TNF (at 200ng/mL, MP6-XT22) was purchased from BioLegend and the antibody to induce TNFR2 trimerization (at 1μg/mL, 80M2) was purchased from Hycult Biotech.

### Cell death analysis

Cells were seeded at a density of 1×10^6^ cells/mL. After treatment, cells were trypsinized, washed once and re-suspended in HBSS containing 1μg/mL propidium iodide (PI) or stained with fixable viability dye (eBioscience). Cells were then assessed for cell death by flow cytometry on a FACS Canto II. Data were analyzed by FlowJo software, version 10.2. Alternatively, cells were directly incubated in the presence of 5μg/ml propidium iodide and assessed for viability by acquiring both phase contrast and red fluorescence images at 2 hour intervals at 10x magnification over 24h using the IncuCyte. Confluency and PI fluorescence were measured and analyzed using the IncuCyte Zoom Software (Version 2016A).

### Cell sorting

To isolate primary macrophages from mouse bone marrow, CD11b^+^F4/80^+^ cells were separated using a FACSMelody 3L machine.

### Caspase activity assay

Cell lysates were treated and lysed in DISC lysis buffer (20mM Tris-HCl pH 7.5, 150mM NaCl, 10% (v/v) glycerol, 1% (v/v) Triton X-100, with protease and phosphatase inhibitors) and incubated with 0.5mM DEVD-AMC. Results were normalized to the protein concentration, that was calculated by BCA (Pierce) according to manufacturer’s instructions.

### Multiplex cytokine analysis

Multiplex cytokine analysis (ProcartaPlex, Thermo Scientific) was performed according to the manufacturer’s instructions and assayed on a Bio-Rad Bioplex machine.

### Western Blotting

Cells were lysed using DISC lysis buffer (20mM Tris-HCl pH 7.5, 150mM NaCl, 10% (v/v) glycerol, 1% (v/v) Triton X-100, with protease and phosphatase inhibitors). The insoluble fraction of the lysate was pelleted by centrifugation and removed. Lysates were boiled and run on 4-12% Bis-Tris Gel NuPAGE using MOPS buffer (Invitrogen). Proteins were then transferred onto PVDF-membrane (0.2μm, Thermo Scientific) using the Trans-Blot^®^ Turbo™ Transfer System (Bio Rad) or the Pierce™ Power Blotter (Thermo Scientific), both according to the manufacturer’s instruction. After blocking with PBST containing 5% (w/v) skim milk, membranes were incubated with the indicated primary antibody in either PBST containing 5% skim milk or 5% (w/v) BSA (Fraction V, Sigma) for at least 1h at RT or overnight at 4°C. Protein level expression was acquired using WesternBright ECL (Advansta) and Amersham Hyperfilm ECL (GE Healthcare).

### qPCR

RNA was isolated using GENEzol Reagent (Geneaid) according to the manufacturer’s instructions. cDNA was produced using MultiScribe™ Reverse Transcriptase and SYBR Green qPCR master mix (Thermo Fisher Scientific) was used for running the qPCR. Melting curves showed that single products were formed. The following primers were used: TNF (5’: CCA CCA CGC TCT TCT GTC TA; 3’: CAC TTG GTG GTT TGC TAC GA); B2M (5’: TGG TGC TTG TCT CAC TGA CC; 3’ CCG TTC TTC AGC ATT TGG AT). Relative standard curve analysis was performed using the housekeeping gene B2M and unstimulated samples were used as a calibrator for fold-change.

### RNA sequencing and analysis

Cells were stimulated with 100ng/mL of TNC-TNF for 2h and RNA was extracted using GENEzol Reagent (Geneaid) and cleaned using PureLink RNA Mini Kit (Invitrogen) according to the manufacturer’s instructions. Libraries were prepared and sequenced at the Functional Genomics Center Zurich (Zurich, Switzerland). RNA-Seq data was processed through a standard workflow (https://github.com/csoneson/rnaseqworkflow), including read mapping against the mouse reference genome (Ensembl_GRCm38.90) using STAR(Dobin et al., 2012) and sorting/indexing with samtools(Li et al., 2009). Isoform-level expression estimation using salmon(Patro et al., 2017) and gene-level differential expression (DE) analysis was performed using edgeR(Robinson et al., 2009) with separate contrasts for each genotype (treated with TNC-TNF versus untreated).

### Statistical analysis

All data is presented in mean ± SEM. Figures were prepared in Illustrator CC 2015 (Adobe) and Prism 7 (GraphPad Software). Significance between genotypes and treatments was assessed by Student-t test or two-way ANOVA with p *=<0.05, **=<0.01, ***=<0.001, ****=<0.0001 using Prism 7.

## Supplementary Figures

**Figure S1.**
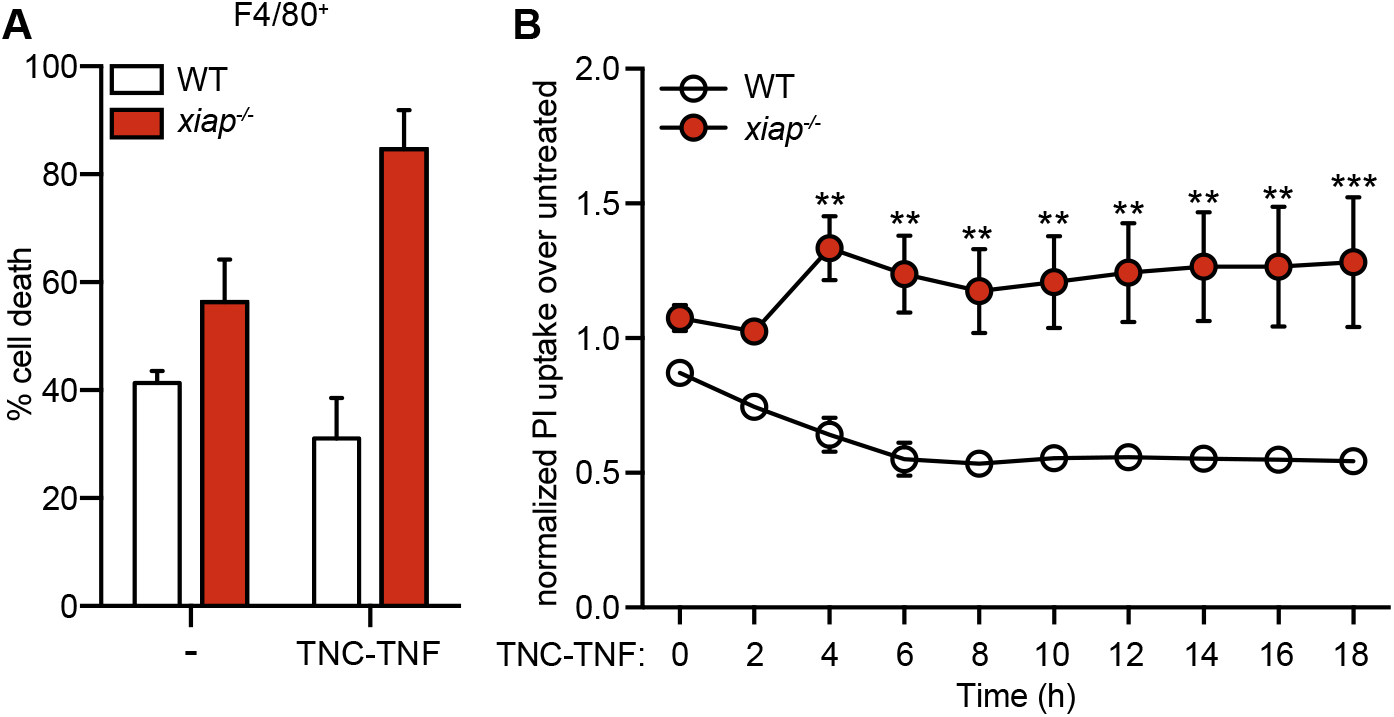
TNFR2-TNF induces cell death in freshly isolated macrophages lacking XIAP. (A) Sorted bone marrow macrophages (CD11b^+^F4/80^+^) from WT and *xiap*^−/−^ mice were treated with TNC-TNF overnight. Cell death was measured via PI uptake by flow cytometry. (B) The course of cell death in WT and *xiap*^−/−^ BMDMs upon TNC-TNF stimulation was monitored over 18h by live imaging using using time-lapse photography (Incucyte). Data shown is mean ± SEM, including n=2-4 biological replicates, experiment was repeated independently for at least three times. Statistical significance was calculated using two-way ANOVA with p **=<0.01, ***=<0.001.

**Figure S2.**
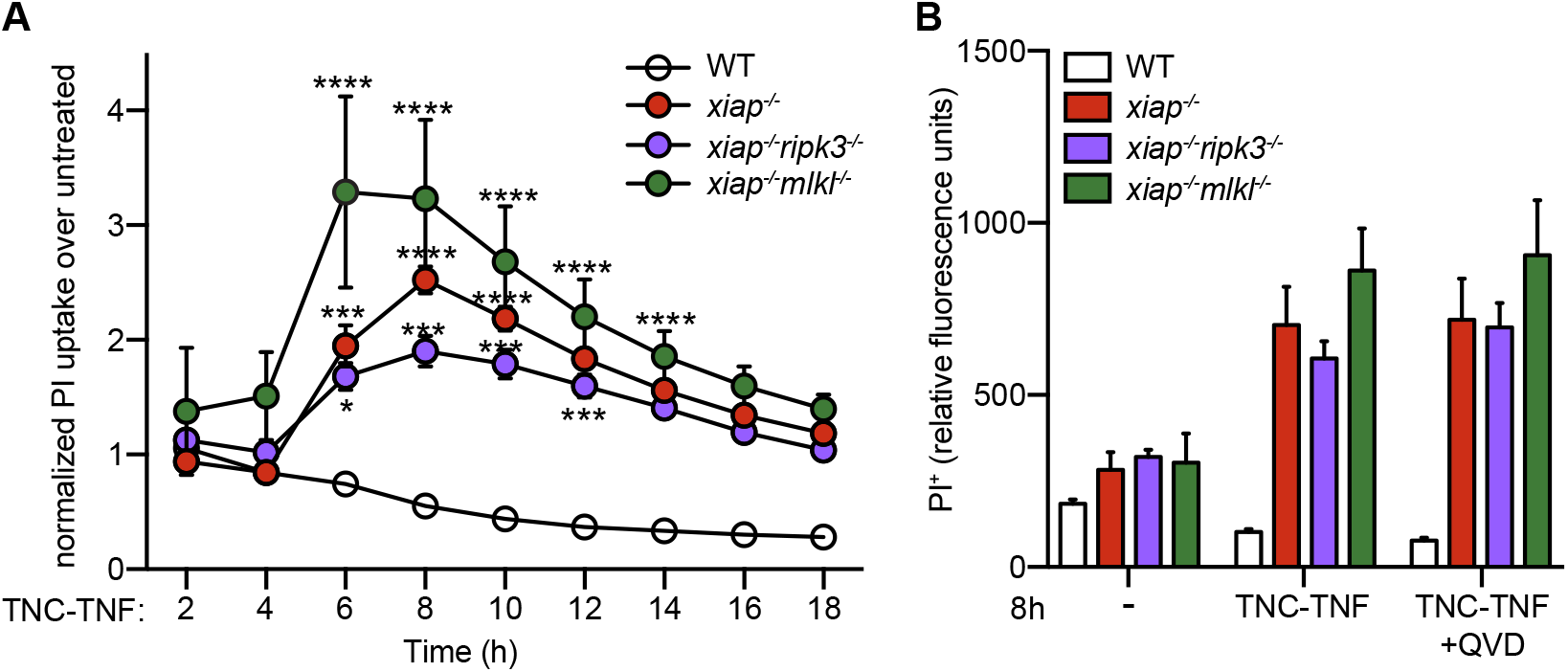
TNFR2-TNF induced cell death in *xiap*^−/−^ BMDMs is independent of necroptosis or caspase activity. (A) WT, *xiap*^−/−^, *xiap*^−/−^*ripk3*^−/−^, *xiap*^−/−^*mlkl*^−/−^ BMDMs were treated with TNC-TNF and monitored over 18 hours for cell death by PI uptake using time-lapse photograph. (B) PI^+^ BMDMs after 8h stimulated with TNC-TNF and/or QVD using time-lapse photography. Data shown is mean ± SEM including n=3 biological replicates. Experiment was repeated three times independently. Statistical significance was calculated using two-way ANOVA with p *=<0.05, **=<0.01, ***=<0.001, ****=<0.0001.

**Figure S3.**
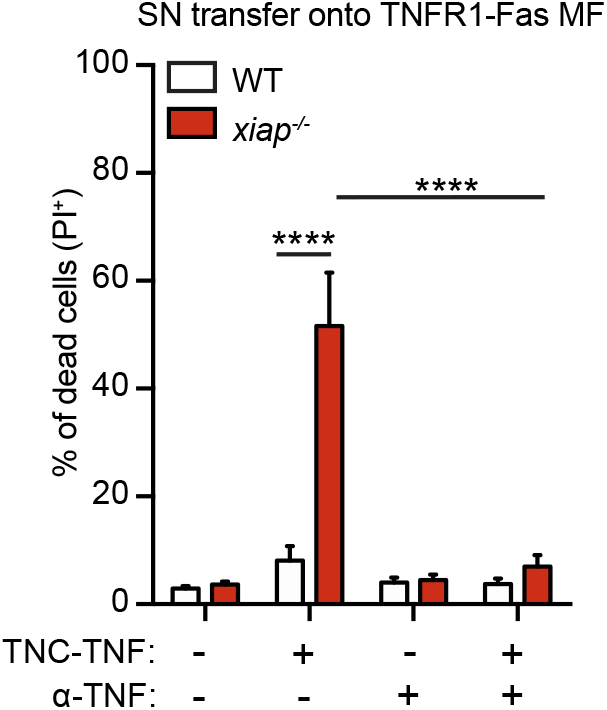
TNFR2 mediated cell death is due to soluble TNF. (A) WT and *xiap*^−/−^ BMDMs were treated with TNC-TNF or in combination with anti-TNFα for 24h. Supernatant was then transferred onto TNFR1-Fas MF cells and after another 24h cell death was measured via PI uptake by flow cytometry. Data is shown as mean ± SEM, including n=5 biological replicates and three independent experiments have been performed. Statistical significance was calculated using two-way ANOVA with p ****=<0.0001.

**Figure S4:**
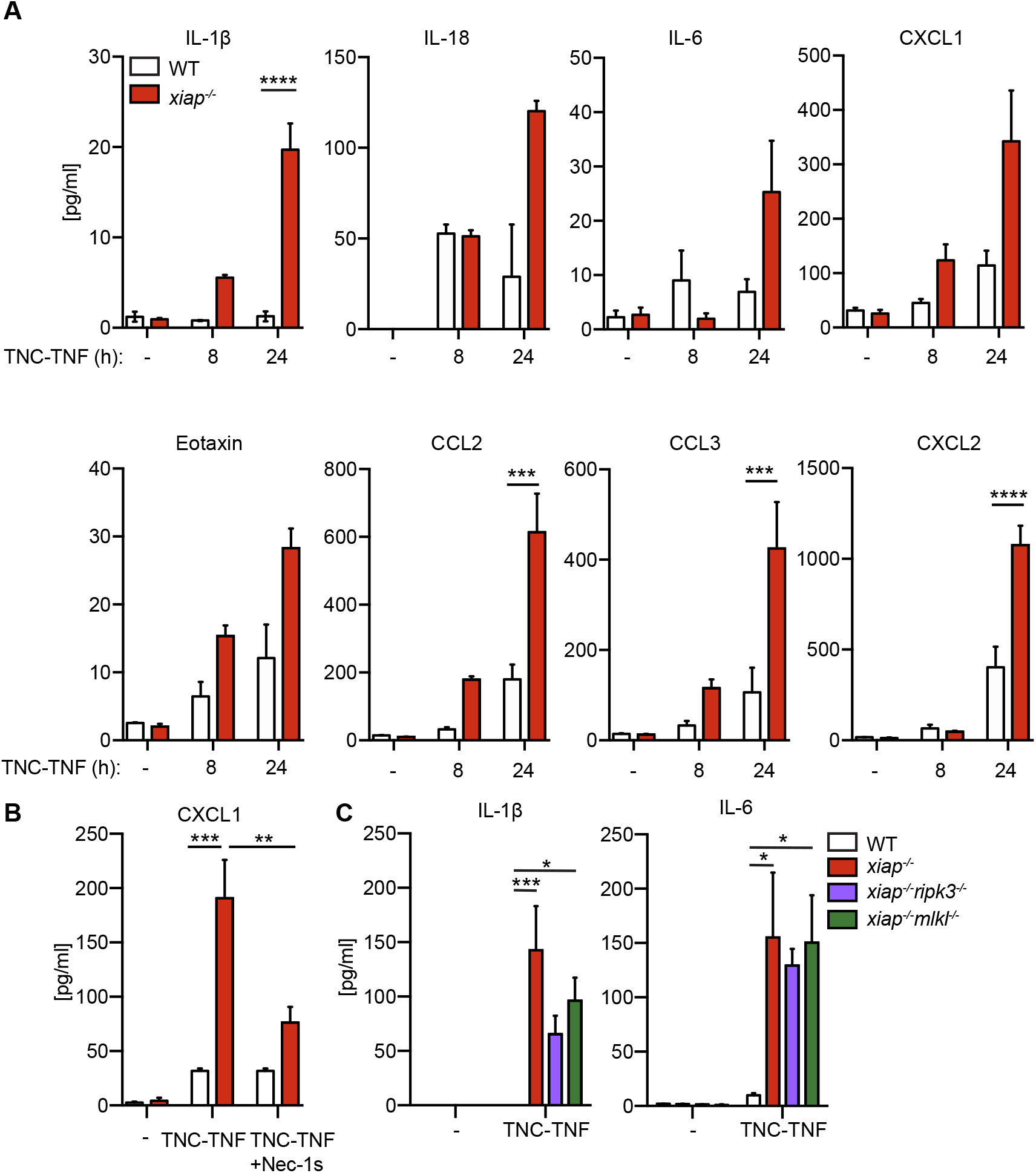
TNFR2 induced cytokine production in *xiap*^−/−^ BMDMs is dependent on RIPK1 kinase activity. WT and *xiap*^−/−^ BMDMs were treated with TNC-TNF. (A) After 0h, 8h and 24h of stimulation supernatant was taken and assayed for IL-1β, IL-18, IL-6, CXCL1, Eotaxin, MIP1α (CCL3), MIP1β (CCL4) and MIP2 (CXCL2) by multiplex. (B) BMDMs were treated with TNC-TNF and/or Nec-1s overnight and supernatants were assayed for CXCL1 by multiplex. (C) BMDMs of indicated genotypes were treated with TNC-TNF for 12h and supernatant was assayed for IL-1β and IL-6. Data shown is mean ± SEM including n=3 biological replicates. Experiment was repeated three times independently. Statistical significance was calculated using two-way ANOVA with p *=<0.05, **=<0.01, ***=<0.001, ****=<0.0001.

**Table S1. RNAseq data of wildtype, *xiap*^−/−^ and *xiap*^−/−^*tnfr1*^−/−^ macrophages in response to TNC-TNF.** BMDMs were treated for 2h with TNC-TNF and transcriptional profiling of the RNA was performed.

**Table S2. Gene set enrichment analysis of TNFR2-induced gene signatures**.

